# M1-linked ubiquitination facilitates NF-κB activation during sterile inflammation

**DOI:** 10.1101/2021.06.03.446895

**Authors:** Anna Aalto, Gabriela Martínez-Chacón, Nadezhda Tsyganova, Joose Kreutzer, Pasi Kallio, Meike Broemer, Annika Meinander

**Affiliations:** Faculty of Science and Engineering, Cell Biology, Åbo Akademi University, BioCity, Turku, Finland; Faculty of Medicine and Health Technology, BioMediTech, Tampere University, Tampere, Finland; German Center for Neurodegenerative Diseases (DZNE), Bonn, Germany

**Author notes:** Corresponding author: Annika Meinander, PhD, Faculty of Science and Engineering, Cell Biology, Åbo Akademi University, Tykistökatu 6A, BioCity 2^nd^ floor, FI-20520 Turku.

**Keywords:** cell stress, hypoxia, linear ubiquitin chain, NF-κB, sterile inflammation

## Abstract

Methionine 1 (M1)-linked ubiquitination plays a key role in the regulation of inflammatory nuclear factor-κB (NF-κB) signalling and is important for clearance of pathogen infection in *Drosophila melanogaster*. M1-linked ubiquitin (M1-Ub) chains are assembled by the linear ubiquitin E3 ligase (LUBEL) in flies. Here, we have studied the role of LUBEL in sterile inflammation induced by different types of cellular stresses. We have found that LUBEL catalyses formation of M1-Ub chains in response to hypoxic, oxidative and mechanical stress conditions. LUBEL is shown to be important for flies to survive low oxygen conditions and paraquat-induced oxidative stress. This protective action seems to be driven by stress-induced activation of the NF-κB transcription factor Relish via the Immune deficiency (Imd) pathway. In addition to LUBEL, the intracellular mediators of Relish activation, including the *Drosophila* inhibitor of apoptosis (IAP) Diap2, the IκB kinase γ (IKKγ) Kenny and the initiator caspase Death-related ced-3/Nedd2-like protein (Dredd), but not the membrane receptor peptidoglycan recognition protein (PGRP)-LC, are shown to be required for sterile inflammatory response and survival. Finally, we showed that the stress-induced upregulation of M1-Ub chains in response to hypoxia, oxidative and mechanical stress is also induced in mammalian cells. Taken together, our results suggest that M1-Ub chains are important for NF-κB signalling in inflammation induced by stress conditions often observed in chronic inflammatory diseases and cancer.

## Introduction

Ubiquitination is a reversible process involving the addition of ubiquitin, a 76-amino acid-long polypeptide, to the target substrate through a three-step enzymatic process carried out by E1 ubiquitin-activating enzymes, E2 ubiquitin-conjugating enzymes and E3 ubiquitin ligases [1]. Polyubiquitin chains are created when a lysine residue (K6, K11, K27, K29, K33, K48, K63) or the N-terminal methionine (M1) on ubiquitin itself are ubiquitinated. These chains are recognised by proteins through ubiquitin-binding domains (UBDs), which translate the ubiquitin modification into cellular outcomes. UBDs can for instance be found in proteins that catalyse ubiquitination and in deubiquitinating enzymes (DUBs) that remove the ubiquitin code from substrates [2–4]. In mammals, M1-linked ubiquitin (M1-Ub) chain formation is catalysed by the linear ubiquitin chain assembly complex (LUBAC) consisting of heme-oxidized iron-responsive element-binding protein 2 ubiquitin ligase-1L (HOIL-1L), HOIL-1-interacting protein (HOIP) and SH3 and multiple ankyrin repeat domains protein (SHANK)-associated RBCK1 homology (RH)-domain-interacting protein (SHARPIN) [5–8]. The role of M1-ubiquitination in inflammatory signalling has been studied extensively in mammals and has been shown to be a key regulator of the canonical nuclear factor-κB (NF-κB) pathway [9–11]. Similarly, in *Drosophila*, the linear ubiquitin E3 ligase (LUBEL)-mediated M1-ubiquitination is induced upon bacterial infection and is required for NF-κB activation, and for mounting a proper immune response in the intestine [12].

In *Drosophila*, infection leads to robust activation of NF-κB signalling pathways, leading to transcription of hundreds of genes, including antimicrobial peptides (AMPs) required for fending off intruding pathogens. During systemic inflammation the expression and release of AMPs is induced in the haemocytes of the fat body, whereas a more local insult leads to an *in situ* production of AMPs. In flies, there are two NF-κB-activating pathways, the Immune deficiency (Imd) pathway and the Toll pathway [13–15]. The Imd pathway is activated upon recognition of Gram-negative bacteria by peptidoglycan recognition proteins (PGRPs), followed by formation of a signalling complex consisting of Imd, the adaptor protein *Drosophila* Fas-associated death domain (dFadd), the caspase-8 homolog Death-related ced-3/Nedd2-like protein (Dredd), and the *Drosophila* Inhibitor of apoptosis protein 2 (Diap2) [16–18]. For transcriptional activation, the NF-κB transcription factor Relish needs to be phosphorylated by the IκB kinase (IKK) complex consisting of the catalytic subunit Ird5 (IKKβ) and the regulatory subunit Kenny (IKKγ) [19–21]. In addition, Relish is cleaved by Dredd, allowing its translocation to the nucleus and the induction of target gene expression [21–23]. The Toll receptor, on the other hand, is activated in response to Gram-positive bacteria and induces activation and translocation of the NF-κB factors Dorsal and Dorsal-related immunity factor (Dif) [15,24].

Recognition of pathogen-associated molecular patterns (PAMPs) by pattern-recognition receptors (PRR) triggers inflammatory NF-κB signalling. However, activation of NF-κB can also be induced in the absence of pathogens, in both *Drosophila* and mammals. Such sterile inflammation may be induced by molecules leaking out from necrotic cells or by materials and compounds that are recognised as danger-associated molecular patterns (DAMP). These molecules can be found in the extracellular or intracellular milieu for example during pathological conditions triggered by mechanical trauma, hypoxia, radiation and chemicals [25,26]. In addition, it has been suggested that cells exposed to stressful conditions release stress-associated molecular patterns (SAMP), which may lead to similar inflammatory responses [27]. Here, we have studied the role of M1-ubiquitination during sterile inflammation induced by mechanical, hypoxic or oxidative stress in *Drosophila* and mammalian cells. Our results show that M1-Ub chains are required for stress-induced activation of NF-κB and survival of flies during hypoxia or oxidative stress. We, hence, suggest that formation of M1-Ub chains is a conserved, common response to different forms of stresses that enables downstream signalling and activation of NF-κB.

## Results

### LUBEL catalyses formation of M1-Ub chains and is required for survival during hypoxia in *Drosophila*

Hypoxia is a condition where the cellular oxygen level decrease, inducing stress responses to protect from damage. To investigate if M1-Ub chains are induced *in vivo* during hypoxia in *Drosophila*, we placed adult fruit flies in specialised hypoxia chambers on a modified MiniHypoxy platform (Fig. 1A) [28], in which the oxygen content is reduced to 5 % within minutes (Fig. 1B). After exposure to hypoxia, we isolated M1-Ub chains from lysates of whole flies with a GST-tagged recombinant tandem ubiquitin-binding entity (TUBE) specific for M1-Ub chains (M1-TUBE). We found that hypoxia induces an increase in M1-Ub chain formation in control flies (Fig. 1C, lanes 1, 2), but not in *LUBEL*^Δ*RBR*^ mutant flies that lack the catalytic RBR domain of LUBEL, responsible for M1 ubiquitination in *Drosophila* (Fig. 1C, lanes 5, 6). A similar effect was observed when the expression of LUBEL was silenced by inducing expression of RNAi transcripts (*LUBEL-RNAi*) using the daughterless-Gal4 (daGal4) driver, which directs ubiquitous expression of UAS-regulated sequences (Fig. 1C, lanes 3, 4). As we have previously shown that LUBEL-mediated M1-Ub chains are induced by ingested bacteria [12], we wanted to exclude the possibility that the observed M1-ubiquitination in response to hypoxia was due to increased receptor-stimulation by resident commensal bacteria. For this purpose, we generated germ-free axenic flies. Similarly, as in conventionally reared flies, an increase in M1-Ub chains was observed in axenic flies during hypoxic conditions (Fig. 1D), indicating that the hypoxia-induced M1-Ub chain formation is not induced by bacteria.

**Figure 1.**
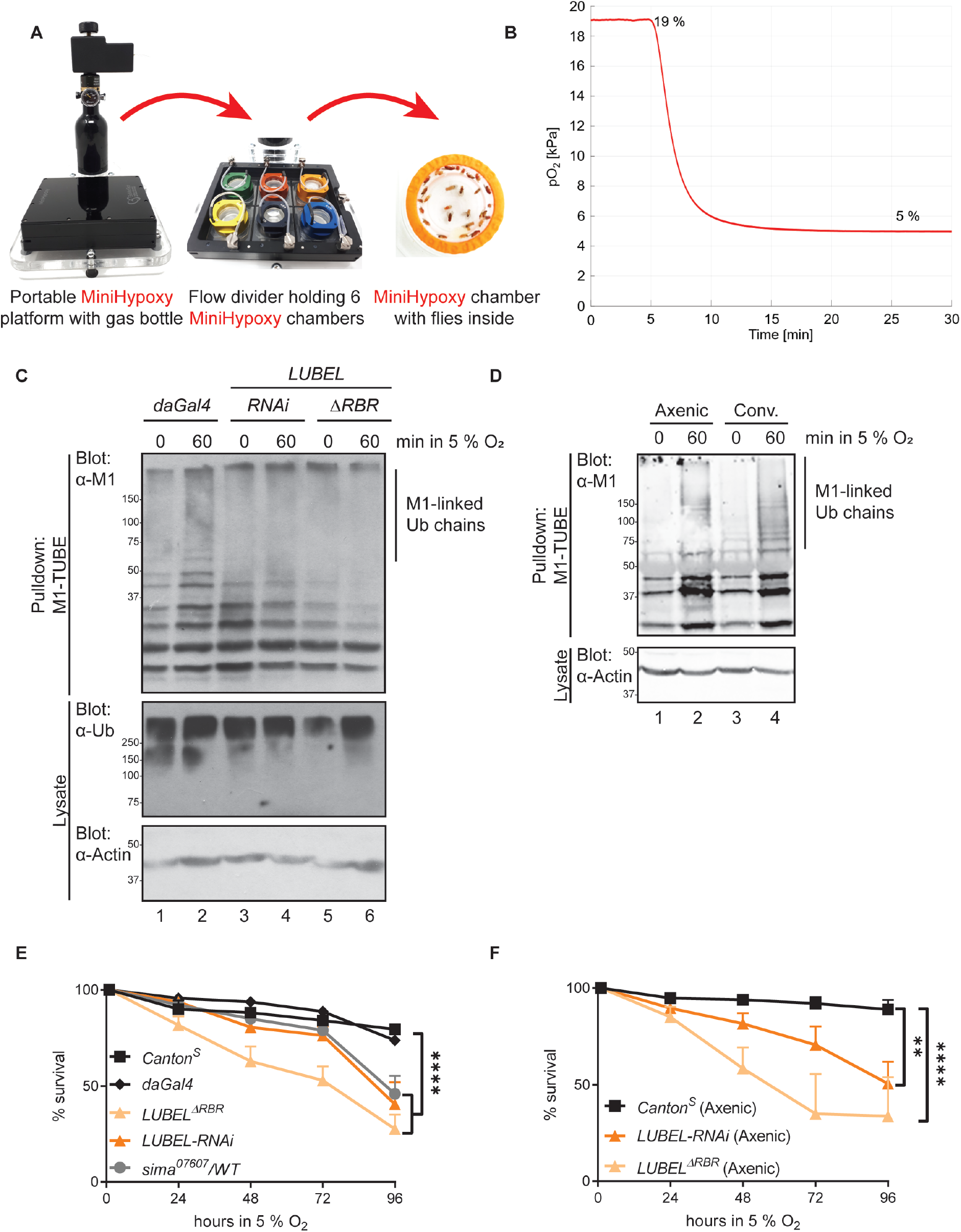
LUBEL catalyses formation of M1-Ub chains and is required for survival during hypoxia in *Drosophila*. **(A)** A picture of the modified MiniHypoxy platform connected to a gas bottle, the 6-well flow divider and a MiniHypoxy chamber with flies. **(B)** The dynamics of gas supply to the MiniHypoxy chamber was monitored by measuring the partial oxygen pressure (pO_2_) while a 5 ml/min gas flow was supplied to the chamber. **(C)** Adult control *daGal4* flies and mutant *LUBEL*^Δ*RBR*^ and *UAS-LUBEL-RNAi;daGal4* flies were subjected to low oxygen conditions (5 % O_2_). M1-Ub chains were isolated at denaturing conditions from fly lysates with M1-TUBE. Ubiquitin chains from samples were analysed by Western blotting with α-M1 and α-pan-Ub antibodies, equal loading was controlled with α-Actin antibody, n=3. **(D)** Conventionally reared control and axenic wild-type *Canton^S^* flies were subjected to low oxygen conditions (5 % O_2_). M1-Ub chains were isolated and analysed as in C, n=3. **(E)** Adult *Canton^S^* and *daGal4* control flies, mutant *LUBEL*^Δ*RBR*^, *UAS-LUBEL-RNAi;daGal4* and heterozygote mutant *sima^K607607^* flies were subjected to low oxygen conditions (5 % O_2_) and their survival was monitored over time. Error bars indicate SEM from more than 10 independent experimental repeats. **(F)** Axenic adult wild-type *Canton^S^* and mutant LUBEL^Δ*RBR*^ and *UAS-LUBEL-RNAi;daGal4* flies were subjected to low oxygen conditions (5 % O_2_) and their survival was monitored over time. Error bars indicate SEM from more than 3 independent experimental repeats. Statistical significance was calculated using two-way ANOVA and Tukey’s multiple comparisons test (E, F), ** p < 0.01, **** p < 0.0001.

To investigate whether LUBEL is needed for *Drosophila* to endure hypoxia, we assessed the survival of LUBEL mutant flies under low oxygen conditions. While wild type flies survived in 5 % O_2_, both *LUBEL*^Δ*RBR*^ and *LUBEL-RNAi* flies were more sensitive to the hypoxic conditions (Fig. 1E). Their sensitivity to hypoxia was similar as in flies lacking one allele of Similar (Sima), the *Drosophila* hypoxia-inducible factor (HIF)-1α [29]. To ascertain that the hypoxia-sensitivity of LUBEL mutant flies was not caused by a defect in maintenance of the commensal bacteriome, we looked at the survival of flies reared axenic. As expected, axenic *LUBEL*^Δ*RBR*^ mutant flies and *LUBEL-RNAi* flies were also more sensitive to hypoxia than wild type axenic flies (Fig. 1F).

### LUBEL is not required for activation of the HIF pathway

The hypoxia-inducible factor HIF is a conserved transcription factor, responsible for activation of expression of genes controlling oxygen homeostasis. To analyse if M1-ubiquitination is needed for activation of HIF-mediated hypoxia responses, we analysed the expression of the hypoxia-inducible Sima target gene *ldh* [30–32] in control and LUBEL mutant flies. The *ldh* expression was induced in control flies, whereas *sima^07607^* mutant flies were unable to induce *ldh* expression upon hypoxia (Fig. 2A). Interestingly, *ldh* was induced equally well in *LUBEL*^Δ*RBR*^ (Fig. 2A), *LUBEL-RNAi* flies (Fig. 2B) and in control flies upon hypoxia. Fatiga is an oxygen sensing hydroxylase that marks Sima for degradation under normal oxygen levels. Fatiga is inactivated in the absence of oxygen, leading to a stabilisation of Sima and a subsequent increase in HIF target gene expression [30,33,34]. Interestingly, the hypoxia-sensitivity of *LUBEL*^Δ*RBR*^ mutants was not rescued by a constitutive activation of the HIF pathway by genetic deletion of Fatiga (Fig. 2C). Similarly, *ldh* expression was induced to comparable levels in control flies and in *LUBEL*^Δ*RBR*^ mutants in the presence and absence of Fatiga (Fig. 2D). These results indicate that LUBEL-mediated M1-Ub chains are neither required for oxygen sensing by Fatiga nor for the activation of HIF.

**Figure 2.**
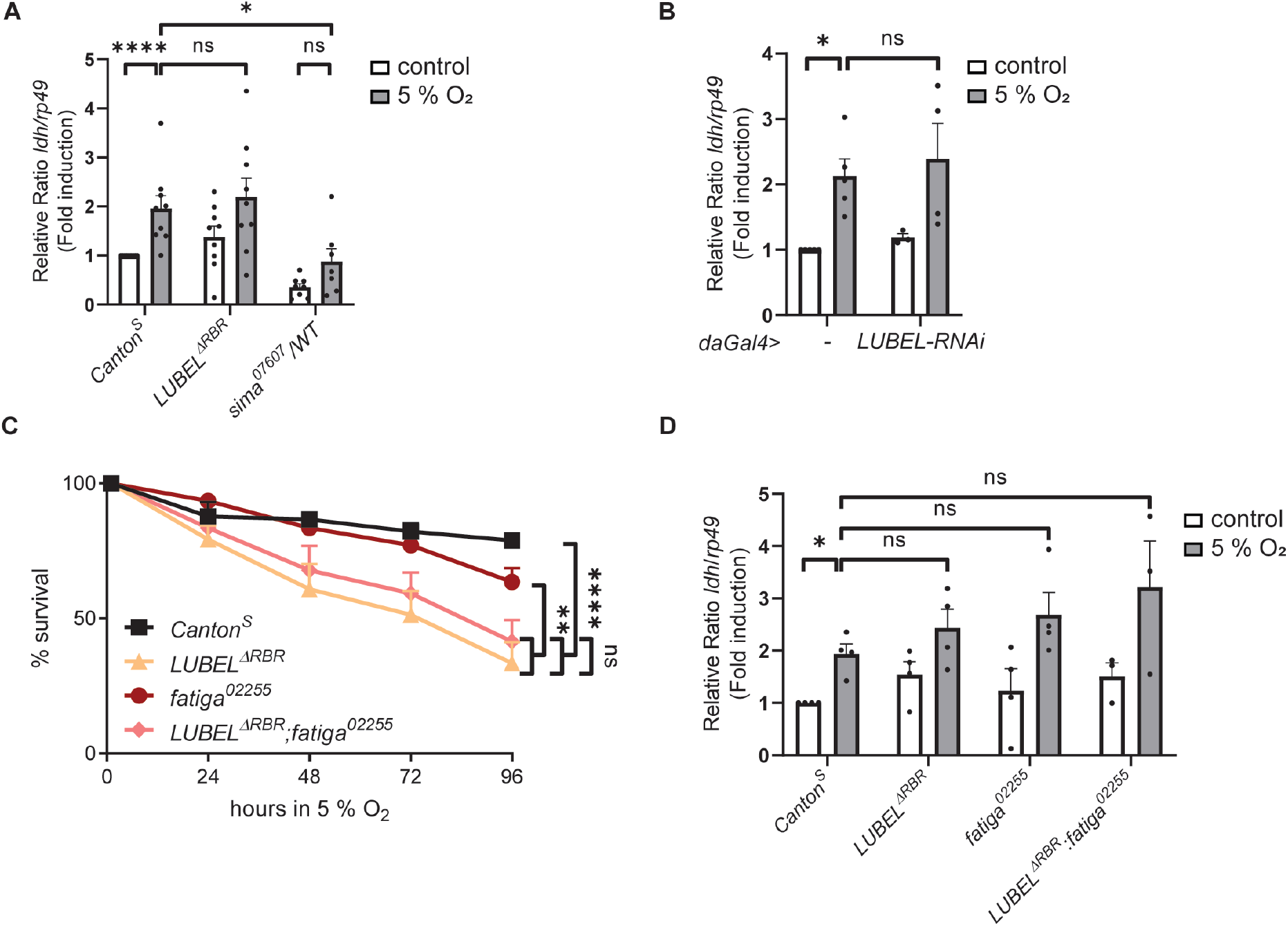
LUBEL is not required for activation of the HIF pathway. **(A, B)** Adult wild type *Canton^S^* or *daGal4* control flies, mutant *LUBEL*^Δ*RBR*^ and *UAS-LUBEL-RNAi;daGal4* flies and heterozygote mutant *sima^K607607^* flies were subjected to low oxygen conditions (5 % O_2_) for 24 h. HIF-pathway activation was studied by analysing the expression of *ldh* with qPCR. Error bars indicate SEM from more than 3 independent experimental repeats. **(C)** Adult wild-type *Canton^S^* flies, mutant *LUBEL*^Δ*RBR*^, mutant *fatiga^02255^* and double-mutant *LUBEL*^Δ*RBR*^; *fatiga^02255^* flies were subjected to low oxygen conditions (5 % O_2_) and their survival was monitored over time. Error bars indicate SEM from more than 6 independent experimental repeats. **(D)** Adult wild-type *Canton^S^* flies, mutant *LUBEL*^Δ*RBR*^, mutant *fatiga^02255^* and double-mutant *LUBEL*^Δ*RBR*^; *fatiga^02255^* flies were subjected to low oxygen conditions (5 % O_2_) for 24 h. HIF-pathway activation was studied by analysing the expression of *ldh* with qPCR. Error bars indicate SEM from more than 3 independent experimental repeats. Statistical significance was calculated using Mann-Whitney U test (A, B, D) or two-way ANOVA and Tukey’s multiple comparisons test (C), ns nonsignificant, * p < 0.05, ** p < 0.01, **** p < 0.0001.

### LUBEL mutant flies are unable to induce Relish target gene expression during hypoxia

As hypoxia has been shown to induce NF-κB activation in flies [32,35], we investigated if this activation requires LUBEL. As expected, expression of the NF-κB target gene *diptericin* was induced during hypoxia in control flies, whereas ubiquitous RNAi silencing of LUBEL prevented the AMP induction (Fig. 3A). *Diptericin* is suggested to be specifically induced by the Imd pathway-specific NF-κB Relish in a Dredd-dependent manner [21,36,37]. To investigate if the LUBEL-mediated induction in *diptericin* expression during hypoxia is mediated via Dredd and Relish, we introduced transgenic expression of Dredd in LUBEL-RNAi flies. Importantly, Dredd overexpression rescued both the inability to induce *diptericin* expression (Fig. 3B) and the sensitivity to hypoxia (Fig. 3C) of *LUBEL-RNAi* flies. Likewise, *LUBEL*^Δ*RBR*^ flies did not die during hypoxia when Dredd expression was induced (Fig. 3D). This indicates that LUBEL regulates hypoxia-induced Relish activation upstream of Dredd.

**Figure 3.**
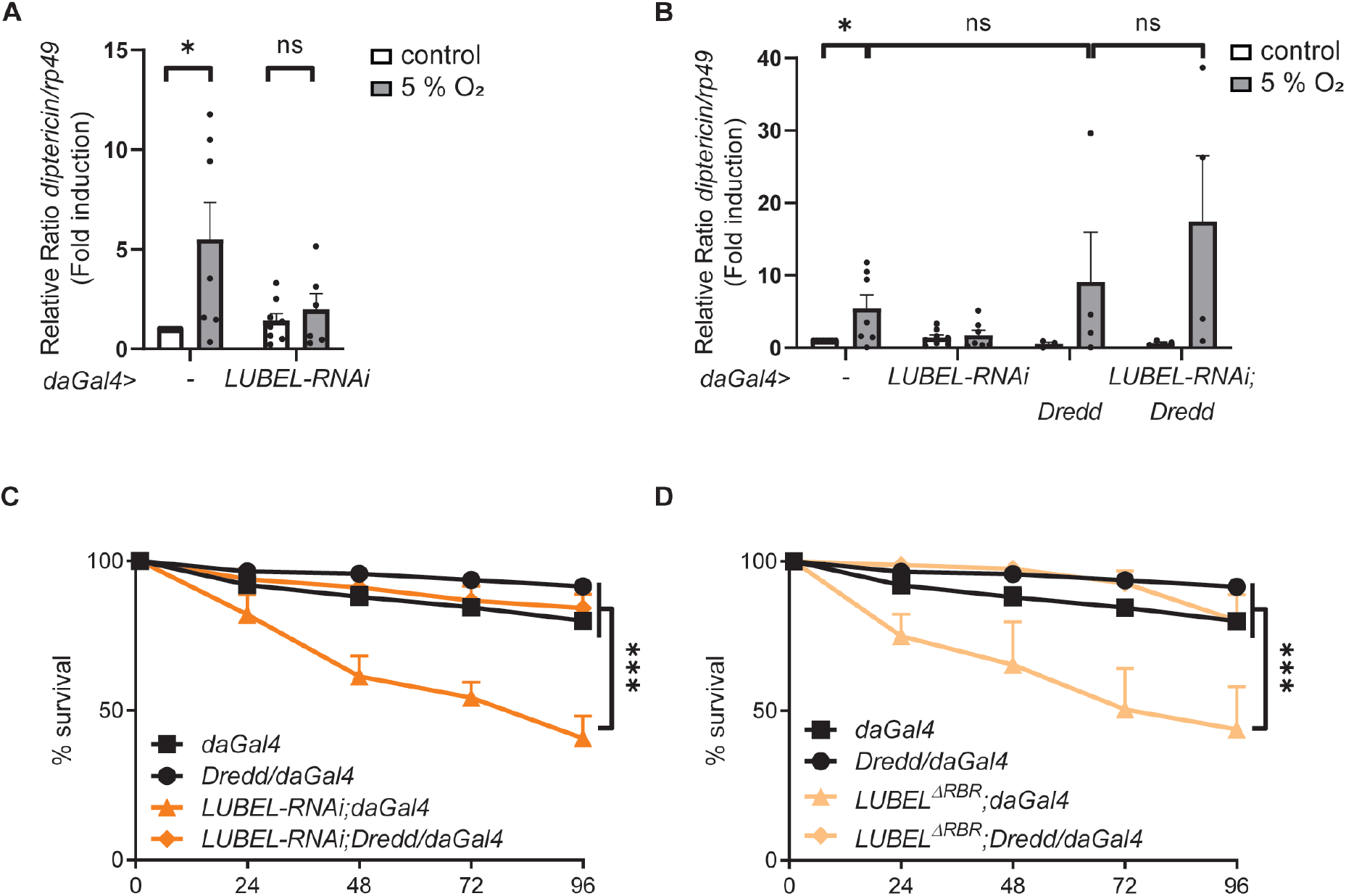
LUBEL mutant flies are unable to induce NF-κB target gene expression during hypoxia. **(A, B)** Adult control *daGal4* flies, mutant *UAS-LUBEL-RNAi;daGal4* flies, overexpression *daGal4/UAS-Dredd* flies and rescue *UAS-LUBEL-RNAi;daGal4/UAS-Dredd* flies were subjected to low oxygen conditions (5 % O_2_) for 24 h. Relish activation was studied by analysing the expression of *diptericin* with qPCR. Error bars indicate SEM from more than 4 independent experimental repeats. **(C, D)** Adult control *daGal4* flies (C, D), overexpressing *daGal4/UAS-Dredd* flies (C, D), mutant *UAS-LUBEL-RNAi;daGal4* (C) and *LUBEL*^Δ*RBR*^ (D) flies, rescue *UAS-LUBEL-RNAi;daGal4/UAS-Dredd* (C) and *LUBEL*^Δ*RBR*^;*daGal4/UAS-Dredd* (D) flies were subjected to low oxygen conditions (5 % O_2_) and their survival was monitored over time. Error bars indicate SEM from more than 4 independent experimental repeats. Statistical significance was calculated using Mann-Whitney U test (A, B) or twoway ANOVA and Tukey’s multiple comparisons test (C, D), ns nonsignificant, * p < 0.05, *** p < 0.001.

### LUBEL is required for both local and systemic AMP expression in response to hypoxia

In flies, NF-κB is induced in the fat body and in epithelial tissues such as the intestine and trachea upon infection [14,38,39]. To study hypoxia-induced NF-κB activation in the trachea, which is the organ for oxygen uptake and distribution, we dissected 3^rd^ instar larvae carrying either a *Drosomycin-LacZ* or *Diptericin-LacZ* reporter. In the trachea, it has been shown that *drosomycin*, but not *diptericin* is expressed upon activation of the Imd pathway [38,40,41]. *Drosomycin* was clearly induced in the trachea of *Drosomycin-LacZ* larvae, but not in *LUBEL*^Δ*RBR*^ mutant *Drosomycin-LacZ* larvae (Fig. 4A). Agreeing with previous findings [38], we could not detect any specific *Diptericin-LacZ* reporter activation in the trachea (Fig. 4B). These data indicate that LUBEL is required for activation of Relish in trachea during hypoxia.

**Figure 4.**
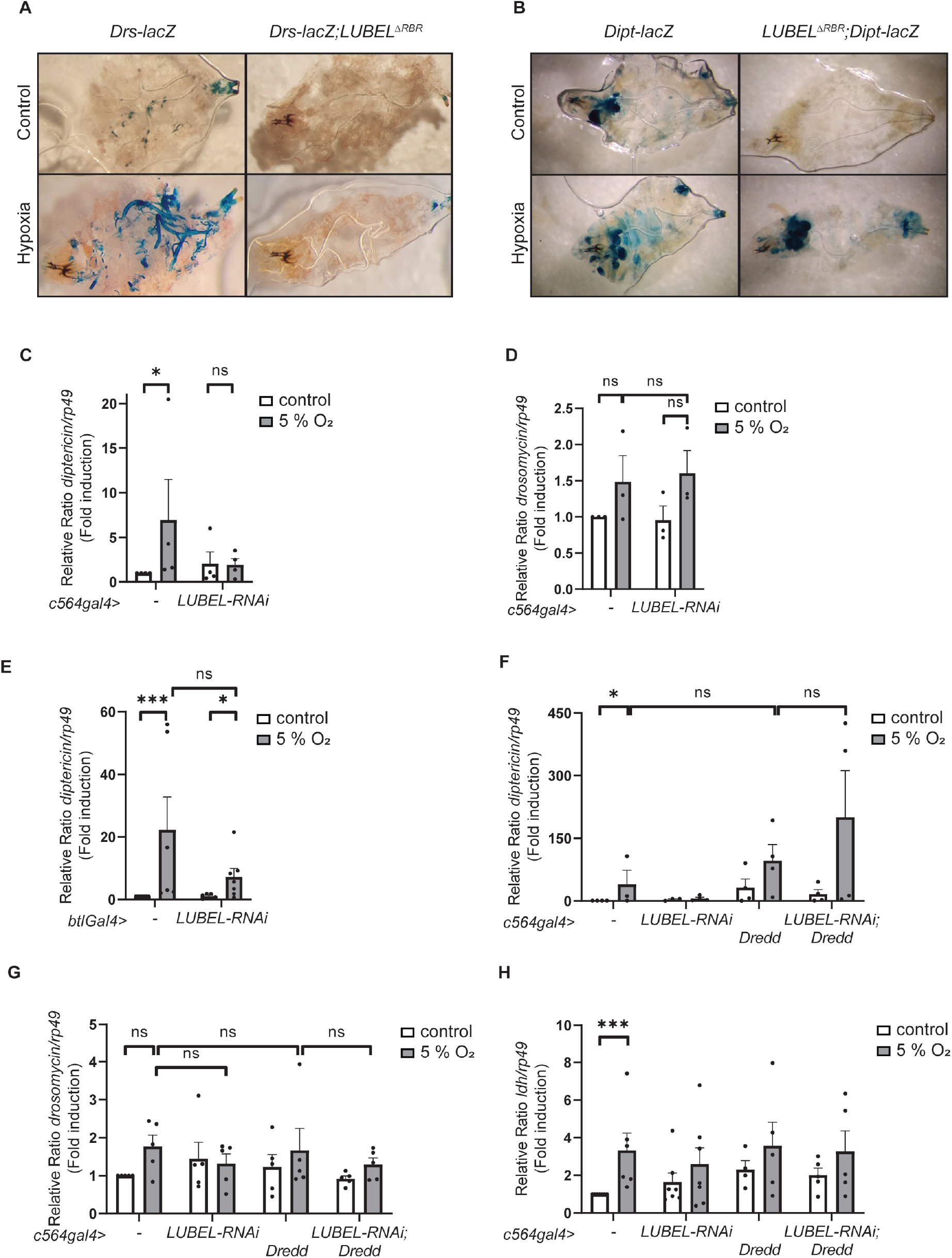
The fly fat body is the responsible for hypoxia-induced NF-κB activation. **(A)** Dissected larval trachea from control *Drosomycin-lacZ* and mutant *Drosomycin-lacZ*;*LUBEL*^Δ*RBR*^ reporter flies stained for β-galactosidase activity after 24 hours of low oxygen treatment (5 % O_2_). The images are representatives of 4 independent experimental repeats. The tracheal structures are visible in the hypoxia-treated control flies in the lower left panel. **(B)** Dissected larval trachea from control *Diptericin-lacZ* and mutant *LUBEL^Δ*RBR*^; Diptericin-lacZ* reporter flies stained for β-galactosidase activity after 24 hours of low oxygen treatment (5 % O_2_). The images are representatives of 3 independent experimental repeats. **(C, D)** Adult control *c564Gal4* flies and mutant *UAS-LUBEL-RNAi/c564Gal4* flies were subjected to low oxygen conditions (5 % O_2_) for 24 h. NF-κB activation was studied by analysing the expression of *diptericin* (C) and *drosomycin* (D) with qPCR. Error bars indicate SEM from more than 3 independent experimental repeats. **(E)** Adult control *btlGal4* flies and mutant *UAS-LUBEL-RNAi/btlGal4* flies were subjected to low oxygen conditions (5 % O_2_) for 24 h. Relish activation was studied by analysing the expression of *diptericin* with qPCR. Error bars indicate SEM from more than 6 independent experimental repeats. **(F)** Adult control *c564Gal4* flies, mutant *UAS-LUBEL-RNAi/c564Gal4* flies, overexpressing *c564Gal4;UAS-Dredd* flies and *UAS-LUBEL-RNAi/c564Gal4;UAS-Dredd* rescue flies were subjected to low oxygen conditions (5 % O_2_) for 24 h. NF-κB activation was studied by analysing the expression of *diptericin* with qPCR. The same samples were analysed for expression of *drosomycin* **(G)** and *ldh* **(H)** with qPCR. Error bars indicate SEM from more than 4 independent experimental repeats. Statistical significance was calculated using Mann-Whitney U test, ns nonsignificant, * p < 0.05, ** p < 0.01, *** p < 0.001.

While we found expression of *diptericin* not to be induced in the trachea, others have shown this AMP to be important for flies to endure hypoxic conditions [32,35]. As *diptericin* is a specific Relish target gene in the fat body [38,42], we analysed if LUBEL regulates Relish activity in the fat body during hypoxia. For this purpose, we specifically silenced LUBEL in the fat body with the fat body-specific driver c564Gal4. While the *diptericin* levels were increased in control flies during hypoxia, no hypoxia-induced *diptericin* expression could be detected when silencing LUBEL in the fat body (Fig. 4C). Meanwhile, *drosomycin* expression, which in contrary to the trachea, is mainly induced via the Toll pathway in the fat body, was not affected by fat body-specific loss of LUBEL (Fig. 4D). This suggests that LUBEL is required for activation of Relish in the fat body during hypoxia. Interestingly, downregulation of LUBEL by driving *LUBEL-RNAi* with the trachea-specific driver btlGal4 did not affect *diptericin* expression in the whole fly (Fig. 4E). These results indicates that while LUBEL affects Relish activation both in the trachea and in the fat body, local Relish activation in the trachea is not needed for systemic Relish activation in the fat body.

To further verify that the LUBEL-mediated activation of *diptericin* gene expression is mediated via Relish, we induced Relish activation by transgenic expression of Dredd in the fat body. Overexpression of Dredd drives *diptericin* expression during hypoxia similarly well in the presence and absence of LUBEL (Fig. 4F), while Dredd is not able to drive expression of *drosomycin* (Fig. 4G) and *ldh* (Fig. 4H). This indicates that LUBEL activates Imd signalling upstream of Dredd and Relish in the fat body. Hence, we conclude that LUBEL is required for both local and systemic Relish activation in response to hypoxia.

### Intracellular but not extracellular mediators are required for Relish activation in response to hypoxia

To further assess the involvement of Relish activation in hypoxia responses, we examined whether other regulators of the Imd pathway are involved. Indeed, the key Imd pathway mediators Dredd, Diap2 and Kenny were all required for flies to survive hypoxic conditions as flies lacking one allele of Kenny, both alleles of Diap2, or carrying a mutation disturbing the catalytic activity of Dredd died during hypoxia (Fig. 5A). Furthermore, Dredd, Diap2 and Kenny mutant flies were unable to induce *diptericin* expression in response to hypoxia (Fig. 5B), while the hypoxia-induced *ldh* expression was comparable in both control and Imd pathway mutant flies (Fig. 5C). Similarly as bacteria do not play a role in the hypoxia-induced M1-ubiquitination and LUBEL-mediated protection from hypoxic conditions (Fig. 1B, D), loss-of-function mutations in the genes encoding the PRRs PGRP-LC and Toll did not induce sensitivity to low oxygen levels (Fig. 5D). This indicates that hypoxia-induced Relish activation is not activated by the conventional PRR but mediated by other intracellular cues.

**Figure 5.**
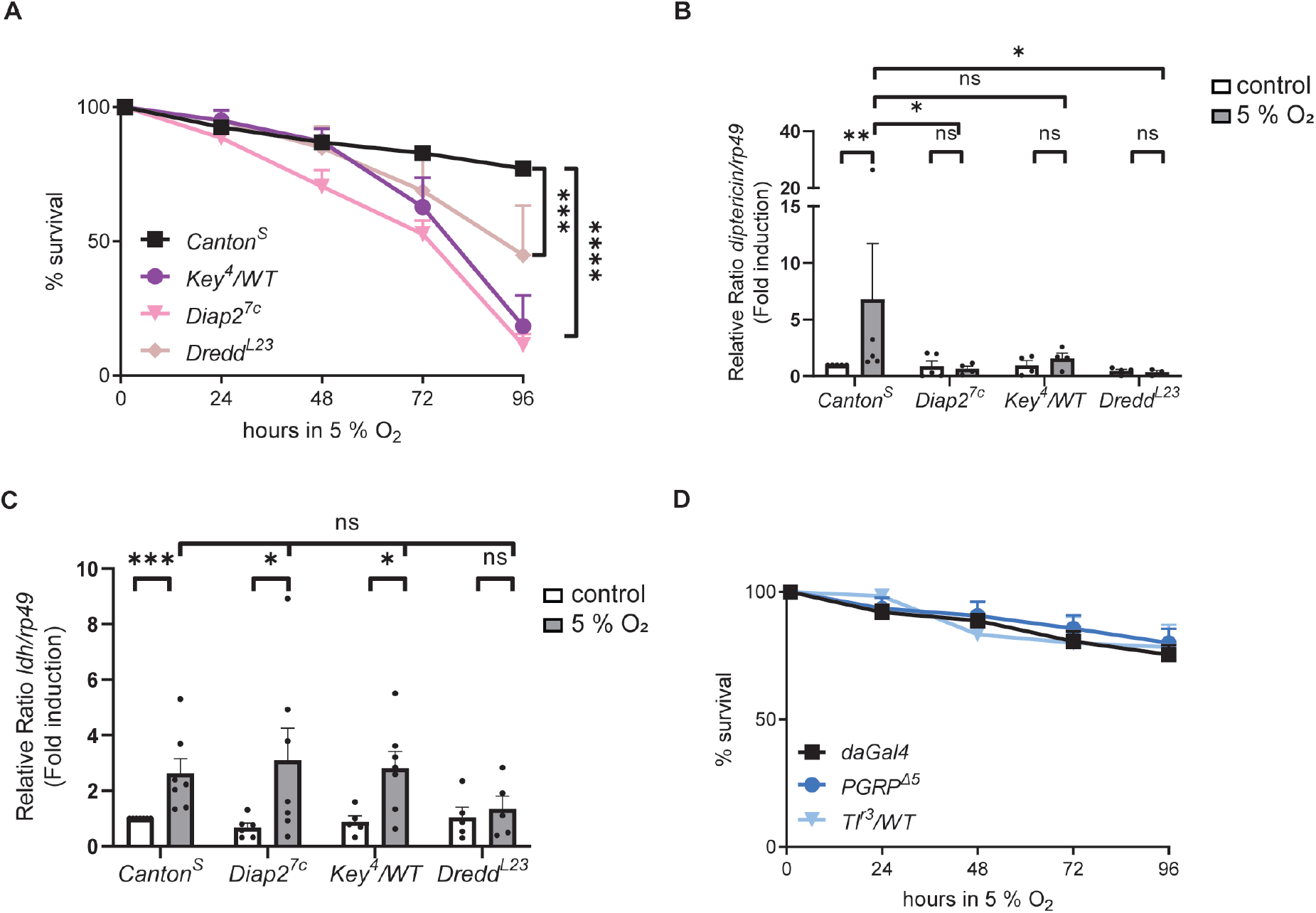
Dredd, Diap2 and Kenny are required for NF-κB activation in response to hypoxia, while the receptors PGRP and Toll are not. **(A)** Adult wild-type *Canton^S^* flies and mutant fly lines of the Imd pathway mediators *Diap2^7c^, Key^4^/WT* and *Dredd^L23^* were subjected to low oxygen conditions (5 % O_2_) and their survival was monitored over time. Error bars indicate SEM from more than 5 independent experimental repeats. **(B, C)** Adult wild-type *Canton^S^* flies and mutant fly lines of the Imd pathway mediators *Diap2^7c^, Key^4^/WT* and *Dredd^L23^* were subjected to low oxygen conditions (5 % O_2_) for 24 h. Relish and HIF activation was studied by analysing the expression of *diptericin* and *ldh*, respectively, with qPCR. Error bars indicate SEM from more than 4 independent experimental repeats. **(D)** Adult control flies *daGal4* and mutant fly lines of the receptors *PGRP*^Δ*5*^ and heterozygote *Tl^r3^/WT* were subjected to low oxygen conditions (5 % O_2_) and their survival was monitored over time. Error bars indicate SEM from more than 3 independent experimental repeats. Statistical significance was calculated using two-way ANOVA and Tukey’s multiple comparisons test (A, D) or Mann-Whitney U test (B, C), ns nonsignificant, * p < 0.05, ** p < 0.01, *** p < 0.001, **** p < 0.0001.

### LUBEL catalyses formation of M1-Ub chains in response to oxidative and mechanical stress

While the Imd pathway did not seem to be induced during hypoxia via receptors recognising extracellular signals, we wanted to address if other types of cellular stresses induce Relish activation, and whether these responses are mediated by LUBEL. To induce oxidative stress, we fed flies with the pesticide paraquat [43,44] and found, interestingly, paraquat feeding was able to induce formation of M1-Ub chains in control flies (Fig. 6A). To elucidate whether LUBEL is important for surviving oxidative stress, we monitored the survival of *LUBEL*^Δ*RBR*^ and *LUBEL-RNAi* flies fed with paraquat during two days. While more than half of control flies survived paraquat feeding, most *LUBEL*^Δ*RBR*^ and *LUBEL-RNAi* flies succumbed (Fig. 6B). The sensitivity was similar to flies in which Dual oxidase (Duox) is silenced by RNAi, rendering the flies unable to produce reactive oxygen species (ROS), thereby, seemingly sensitising them to oxidative stress (Fig. 6B). As during hypoxia, ectopic expression of Dredd was able to rescue the stress-sensitive phenotype of the *LUBEL*^Δ*RBR*^ (Fig. 6C) and *LUBEL-RNAi* flies (Fig. 6D). This suggests that LUBEL induces Imd signalling in response to oxidative stress. Loss-of-function of the receptors PGRP-LC and Toll did not affect sensitivity to paraquat feeding (Fig. 6E), indicating that these NF-κB activating receptors are also not required for protection against oxidative stress. As mechanical stress has been shown to activate NF-κB in *Drosophila* [45], we also investigated if M1-Ub chains are induced in response to mechanical shear stress. We induced shear stress in 3^rd^ instar larvae by vortexing and were able to detect an increase in M1-Ub chain formation (Fig. 6F). These results indicate that M1-ubiquitination is induced as a response to several stress conditions in the fly, including hypoxia, oxidative and mechanical stress.

**Figure 6.**
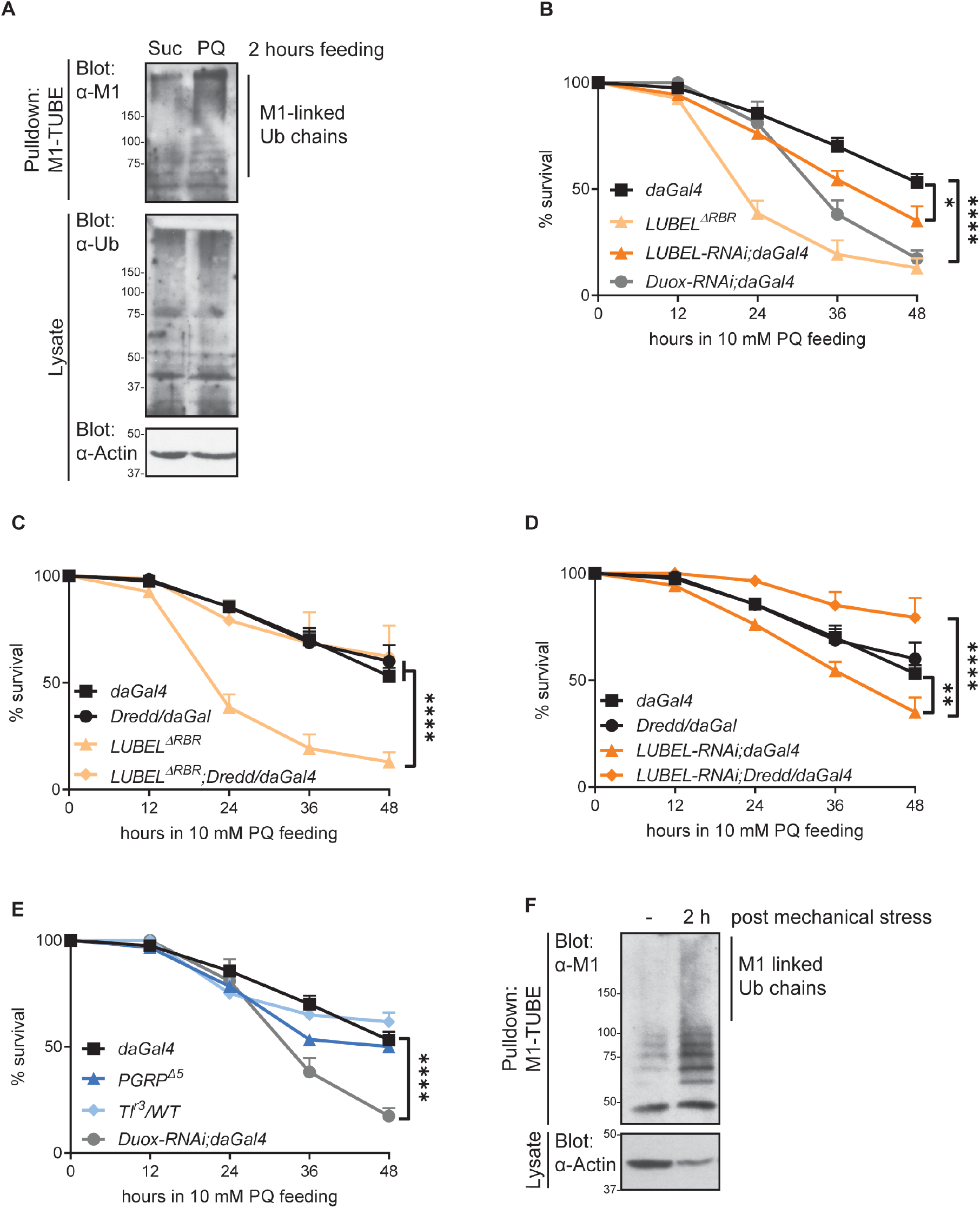
LUBEL catalyses formation of M1-Ub chains in response to oxidative and mechanical stress in *Drosophila*. **(A)** Adult control flies *daGal4* were fed with 5 % sucrose (Suc) or 5 % sucrose with 20 mM paraquat (PQ) for 2 h. M1-Ub chains were isolated at non-denaturing conditions from fly lysates with M1-TUBE. Ubiquitin chains from samples were analysed by Western blotting with α-M1 and α-pan-Ub antibodies, equal loading was controlled with α-Actin antibody, n=3. **(B-E)** Adult control *daGal4* flies (B-E), mutant *LUBEL*^Δ*RBR*^ (B, C), *UAS-LUBEL-RNAi;daGal4* (B, D), *UAS-Duox-RNAi;daGal4* (B, E), overexpressing *daGal4/UAS-Dredd* (C, D), rescue *LUBEL*^Δ*RBR*^;*daGal4/UAS-Dredd* (C), *UAS-LUBEL-RNAi;daGal4/UAS-Dredd* (D), *PGRP*^Δ*5*^ (E) and heterozygote mutant *Tl^r3^/WT* (E) flies were fed with 10 mM paraquat in 5 % sucrose and their survival was monitored over time. Error bars indicate SEM from more than 3 independent experimental repeats. **(F)** Wild-type *Canton^S^* larvae were vortexed for 10 seconds and then placed in normal food to recover for indicated time. M1-Ub chains were isolated at denaturing conditions from fly lysates with M1-TUBE. Ubiquitin chains from samples were analysed by Western blotting with α-M1 antibody, equal loading was controlled with α-Actin antibody, n=3. Statistical significance was calculated using two-way ANOVA and Tukey’s multiple comparisons test, * p < 0.05, ** p < 0.01, **** p < 0.0001.

### M1-Ub chain formation in response to hypoxic, oxidative and mechanical stress is conserved in human intestinal epithelial cells

All the stress conditions that we studied in *Drosophila* are known to induce evolutionary conserved stress-induced inflammatory responses. To investigate if the augmentation of M1-Ub chain formation is an evolutionary conserved response in mammals, we exposed human cancerous intestinal epithelial Caco2 cells to the same stress conditions used in this study in flies. Accordingly, Caco2 cells were placed in a hypoxia chamber with 5 % O_2_ to expose cells to hypoxic conditions, treated with paraquat to induce oxidative stress, and shaken on an orbital shaker to induce mechanical shear stress. Interestingly, all these stress conditions led to an increase of M1-Ub chains compared to those in basal state in Caco2 cells (Fig. 7 A, B, C). This indicates that the signalling sensors and mediators through which M1-Ub chains are induced during these stress conditions are conserved also in mammals.

**Figure 7.**
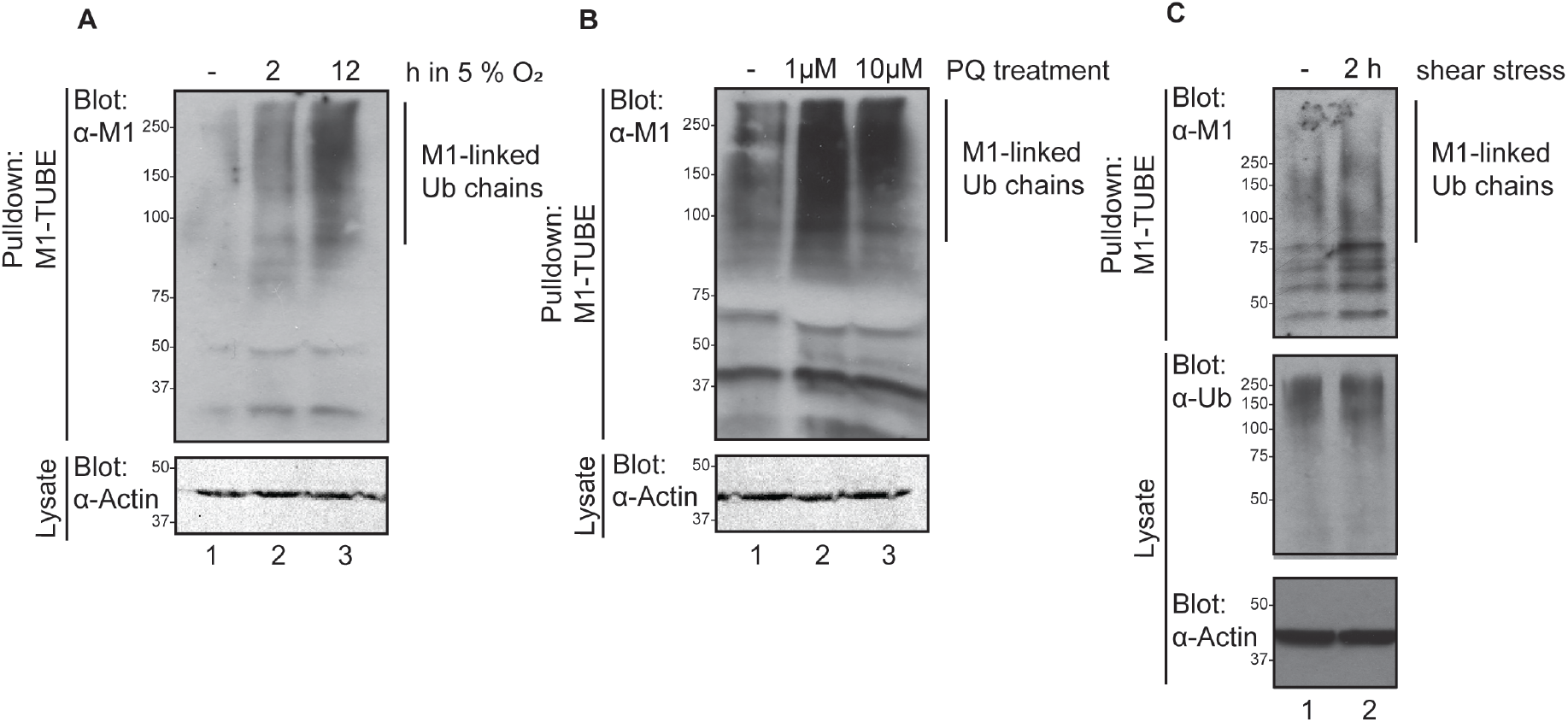
M1-Ub chain formation is induced in response to hypoxic, oxidative and mechanical stress in human intestinal epithelial cells. Human Caco2 cells were **(A)** exposed to low oxygen conditions (5 % O_2_), **(B)** treated 24 h with indicated concentrations of paraquat or **(C)** subjected to shear stress by placing the plates on an orbital shaker, 100 rpm, for 2 h. M1-Ub chains were isolated at denaturing conditions from cell lysates with M1-TUBE. Ubiquitin chains from samples were analysed by Western blotting with α-M1 and α-pan-Ub antibodies, equal loading was controlled with α-Actin antibody, n=3.

## Discussion

During pathological conditions, specific stress responses are induced to provide the cell with the tools to repair or eliminate damaged proteins. In addition, activation of inflammatory signalling cascades, such as the NF-κB pathways contribute to restoration of cellular homeostasis. In the absence of pathogens, this aseptic induction of inflammatory signalling is referred to as sterile inflammation. In this study, we have shown that M1-linked ubiquitination is augmented in response to noxious stimuli like hypoxia and oxidative or mechanical stress, and furthermore that this LUBEL-mediated M1-ubiquitination is required for sterile activation of NF-κB and for *Drosophila* to survive hypoxic and oxidative stress conditions. Importantly, we also demonstrate that M1-Ub chain formation is induced in response to sterile stresses in human cells, indicating that the role of M1-ubiquitination in protection during stress conditions is conserved. We have previously shown that M1-ubiquitination is strongly induced both during septic and local infection in *Drosophila*, but that the M1-Ub chains are indispensable only for mounting local immune responses in the epithelial tissues, and not for NF-κB activation in the fat body [12]. Surprisingly, we have found that LUBEL-mediated M1-ubiquitination in the fat body mediates hypoxia-induced NF-κB activation and may protect against hypoxic conditions. Similarly, it has been shown that NF-κB target gene expression is activated in the fat body during sterile inflammation induced by mechanical pinching of *Drosophila* larvae [45].

Cell stress leads to loss of appropriate folding and protein aggregation, which is recognised by chaperones and leads to ubiquitin-marking of aberrant protein structures [46]. LUBAC has been shown to be recruited to bacteria marked by ubiquitin by direct interaction with the ubiquitin-binding Npl4 zinc finger (NZF) of HOIP [46]. HOIP can also be recruited to ubiquitin-marked molecules indirectly by its peptide N-glycanase (PNG)/UBA or UBX-containing protein (PUB) domain interacting with the PUB-interacting motif (PIM) of the ubiquitin-binding chaperone Valosin-containing protein (VCP) /p97 [47–49]. While stress-induced M1-ubiquitination is a conserved response, also the stress-sensing mechanisms may be conserved. *Drosophila* LUBEL contains several NZF motives in its N-terminus [12,50] that could potentially recognise ubiquitinated molecules during stress. However, an interaction with the VCP/p97 homologue, transitional endoplasmic reticulum 94 (TER94) would most probably require a PUB-containing protein such as Tamo, as no PUB domain is found in the *Drosophila* LUBEL. On the other hand, we cannot completely exclude the role of Toll as an initiator of stress-induced activation of NF-κB, as our studies are performed in heterozygote Toll mutant flies, where a remaining allele may contribute to the functional stress responses. However, we found the PRR receptor PGRP-LC not to mediate Imd pathway activation during stress. Hence, while PAMPs, DAMPs and SAMPs may be patterns of extracellular origin, we hypothesise that the stress-induced activation of M1-ubiquitination is caused by recognition of changes in intracellular patterns.

It is still unknown how cell stress is recognised and converted into activation of inflammatory transcription factors and what the physiological relevance of stress-induced activation of immune responses is in both flies and mammals. Sterile immune responses may function to prime cells to respond to pathogenic invasion, to maintain cellular homeostasis or to support stress-specific protective responses by inducing expression of additional survival-promoting genes. As stress-activated inducers of survival-promoting NF-κB activation, M1-Ub chains may be important protectors of cells that are damaged by protein aggregation, for example in neurodegenerative disorders. On the other hand, M1-Ub chains could contribute to cancer development by protecting cells growing in hypoxic conditions, a state common for growing tumours. M1-Ub chain induction may also protect cancerous cells from mechanical stress caused by changes in cell density and cell-cell adhesion during cancer cell growth and metastasis. In addition, M1-Ub chains might contribute to drug-resistance by activating protective measures in response to genotoxic and proteotoxic stress that is induced by anticancer drugs and by components released from cells killed by cytotoxic agents. Finally, M1-Ub chains may contribute to inflammation-induced cancer by maintaining NF-κB-mediated expression of growth-promoting and anti-apoptotic genes. The highly editable ubiquitin and its regulators serves as interesting targets when tuning inflammatory diseases and cancer. However, further examination of the role of M1-ubiquitination during these pathological conditions is needed to unravel the details of ubiquitin-mediated induction of sterile inflammation.

## Materials and Methods

### Fly husbandry and strains

*Drosophila melanogaster* were maintained at 25°C with a 12 h light–dark cycle on Nutri-fly BF (Dutscher Scientific, Essex, UK). *Canton^S^* wild type flies, the *daGal4* driver line, the fat body-specific *c564Gal4* driver line, the *Dipt-LacZ* reporter line, balancer lines, as well as *key^4^, dredd^L23^* and *diap2^7c^* mutant fly lines were kindly provided by Prof. Pascal Meier and Dr. François Leulier [51]. The *Drosophila* fly lines *yw;Mi{ET1}LUBEL^MB00197^* (#22725, referred to as *LUBEL*^Δ*RBR*^), *fatiga^02255^* (#11561), *sima^KG07607^* (#14640), trachea-specific *btlGal4,GFP* driver line (#8807), *PGRP*^Δ*5*^ (#36323), *UAS-Duox-TRiP* (#33975, referred to as *DUOX-RNAi*) and *Drs-lacZ* (#55708) were obtained from Bloomington stock centre. *UAS-LUBEL-RNAi (P{GD7269}v18055*, #18055, referred to as *LUBEL-RNAi*) were obtained from Vienna *Drosophila* resource center. *Tl^r3^* (#106934) was obtained from KYOTO Stock Center (DGRC) in Kyoto Institute of Technology.

### Stress treatments and survival experiments in *Drosophila*

Hypoxia experiments were performed by placing adult flies or larvae in a modified portable MiniHypoxy-platform (Faculty of Medicine and Health Technology, Tampere University, Finland) [28]. The modified MiniHypoxy-platform can hold six individual MiniHypoxy chambers and a single chamber can hold up to 80 flies (Fig. 1A). With the modified portable MiniHypoxy-platform, flies can be treated with a desired gas-mixture, it enables, for example, live monitoring of flies under a microscope throughout an experiment. In this study, flies were exposed to a gas-mixture of 5 % O_2_ with 95 % N_2_. The readymade gas-mixture is supplied either directly from a large cylinder obtained from Linde Gas (Oy Linde Gas Ab, Espoo, Finland) or as in this study, to support portability, from a small refillable gas bottle (Fig. 1A). The refillable gas bottles can be loaded with any gas composition and thus flies can be exposed and maintained in various gas environments. 5 ml/min gas flow was supplied to each chamber, which renders the chambers hypoxic within minutes (with 3 min 20 s fall time) (Fig. 1B). To validate the functionality of the platform, the partial oxygen pressure (pO_2_) was measured using an in-house made optical oxygen sensor [52]. In short, oxygen sensitive fluorescent dyes embedded in thin polymer film on a glass plate at the bottom of the chamber provides a non-invasive way to measure oxygen partial pressure in the chamber through the glass. The establishment of the correct oxygen level in the chamber and the gas exchange dynamics were demonstrated using readymade mixtures of 19 % and 5 % oxygen. For isolation of M1-Ub chains, 40 adult flies per genotype were incubated for 1 h and for qPCR 10 flies per genotype were incubated 24 h at 25°C in the MiniHypoxy chambers with 5 % oxygen. For X-Gal staining larvae were exposed to 5 % oxygen for 24 h before dissection at 25°C. For hypoxia survival assays, 20 flies per fly genotype were counted at indicated time points. To induce oxidative stress, adult flies were fed 20 mM paraquat in 5 % sucrose pipetted on a Whatman paper. For survival assays, more than 10 flies per fly genotype were fed with 10 mM paraquat and counted at indicated time points. For mechanical stress, 3^rd^ instar larvae were subjected to mechanical stress by vortexing for 10 seconds at 3,200 rpm. For isolation of M1-Ub chains, 15 larvae were frozen at indicated time points after vortexing.

### Cell culture and stress treatments of human Caco2 cells

Human epithelial colon adenocarcinoma, Caco2, cells (ACC 169, DSMZ, Leipzig, Germany) were grown in DMEM/F-12 (Gibco, Thermo Scientific, Waltham, Massachusetts, USA) supplemented with 10 % heat inactivated fetal bovine serum (Biowest, Nuaillé, France), 100 IU/ml penicillin and 100 μg/ml streptomycin (Sigma-Aldrich, Missouri, USA) at 37°C humidified atmosphere with 5 % CO_2_ until use. For hypoxia and oxidative stress experiments, cells were plated at 1×10^6^ cells on 10 cm diameter dishes with serum free DMEM/F-12 supplemented with 0,1 % bovine serum albumin (BSA, Sigma-Aldrich) and 100 IU/ml penicillin and 100 μg/ml streptomycin. Hypoxic conditions were achieved by exposing 80-90 % confluent cells to 5 % O_2_, 5 % CO_2_ and 90 % pure N_2_ (AGA, Finland) by placing the plates in a hypoxic incubator (InvivO_2_, Ruskinn Technology, Bridgend, UK) for 2 and 12 h at 37°C. Oxidative stress was induced by treating cells with 1 μM and 10 μM paraquat (Sigma-Aldrich) for 24 h. For the shear stress experiments, Caco2 cells were seeded in 6-well plates. The flow setup for inducing shear stress was prepared as previously described [53] with minor alterations. Briefly, when 100 % confluency of Caco2 cells was obtained, they were subjected to shear stress by placing the plates on an orbital shaker, 100 rpm, for maximum 2 h at 37°C. Only the outer part of the well was collected to ensure a flow with less oscillatory shear index. The exclusion of the inner part was based on measurements done previously [53]. After exposure, cells were lysed for M1-TUBE pulldown assay.

### Plasmids and antibodies

M1-Ub conjugates were purified using a recombinant protein containing the UBAN region of NEMO (residues 257-346) fused to GST and His (referred to as M1-TUBE) [54,55] provided by Dr. Mads Gyrd-Hansen. The following antibodies were used: α-M1 (clone IE3, #MABS199, Millipore, Burlington, Massachusetts, USA or clone 1F11/3F5/Y102L, Genentech, South San Francisco, California, USA), α-pan-Ub (P4D1, #sc8017, Santa Cruz Biotechnology, Dallas, Texas, USA) and α-Actin (C-11, sc-1615, Santa Cruz Biotechnology).

### Purification of His/GST-tagged M1-TUBE fusion protein

His/GST-tagged M1-TUBE expression was induced in *E. coli* BL21 (Novagen/ Merck Millipore, Burlington, Massachusetts, USA) by the addition of 0.2 mM IPTG to an overnight culture of bacteria in LB medium at 18°C. Bacteria were lysed by sonication in lysis buffer containing 50 mM Tris-HCl (pH 7.4), 150 mM NaCl, 2 mM β-mercaptoethanol (Sigma-Aldrich), 40 mM imidazole, Pierce™ Protease inhibitor and 3 mg lysozyme. Clarified lysate was added to a column with Ni-NTA agarose (QIAGEN, Hilden, Germany) and then washed with cold lysis buffer. Bound protein was eluted with cold lysis buffer supplemented with 300 mM imidazole (pH 7.4) and dialysed into PBS (Medicago, Uppsala, Sweden) containing 10 % glycerol and 1 mM DTT (Sigma-Aldrich) using a 10 K Slide-A-Lyzer dialysis cassette (Thermo Scientific).

### Purification of linear ubiquitin conjugates from flies and cells

Flies and cells were lysed using a buffer containing 50 mM Tris pH 7.5, 150 mM NaCl, 1 % TritonX, 1 mM EDTA, 10 % glycerol supplemented with 1 mM DTT, 5 mM NEM, 1 x Pierce™ Protease and phosphatase Inhibitor, 5 mM chloroacetamide and 1 % SDS. Lysates were sonicated, diluted to 0.1 % SDS, and cleared before incubation with Glutathione Sepharose™ 4B (GE Healthcare) and M1-TUBE (30-100 mg/ml) for minimum of 2 h or o/n under rotation at 4°C. The beads were washed 3 times with ice cold wash buffer containing 10 mM Tris pH 7.5, 150 mM NaCl, 0.1 % TritonX, 5 % glycerol and eluted using Laemmli sample buffer.

### Quantitative RT-PCR (qPCR)

Flies were homogenised using QIAshredder (QIAGEN) and total RNA was extracted with RNeasy Mini Kit (QIAGEN) and cDNA was synthesised with SensiFast cDNA synthesis kit (Bioline, London, UK) according to the manufacturers’ protocols. qPCR was performed using SensiFast SYBR Hi-ROX qPCR kit (BioLine). *rp49* was used as a housekeeping gene for ΔΔCt calculations. The following gene-specific primers were used to amplify cDNA: *Diptericin* (5’-ACCGCAGTACCCACTCAATC-3’, 5’-ACTTTCCAGCTCGGTTCTGA-3’), *Drosomycin* (5’-CGTGAGAACCTTTTCCAATATGATG-3’, 5’-TCCCAGGACCACCAGCAT-3’), *ldh* (5’-CAGTTCGCAACGAACGCGCA, 5’-CAGCTCGCCCTGCAGCTTGT-3’), *rp49* (5’-GACGCTTCAAGGGACAGTATCTG-3’, 5’-AAACGCGGTTCTGCATGAG-3’).

### X-gal staining of *Drosophila* trachea

Trachea from 3^rd^ instar fly larvae were dissected in PBS and fixed for 15 minutes with PBS containing 0.4 % glutaraldehyde (Sigma-Aldrich) and 1 mM MgCl_2_ (Sigma-Aldrich). The samples were washed with PBS and incubated with a freshly prepared staining solution containing 5 mg/ml X-gal (5-Bromo-4-chloro-3-indolyl-β-D-galactopyranoside), 5 mM potassium ferrocyanide trihydrate (Sigma-Aldrich), 5 mM potassium ferrocyanide crystalline (Sigma-Aldrich) and 2 mM MgCl_2_ in PBS at 37°C. After washing with PBS, the samples were mounted using Mowiol (Sigma) and imaged with brightfield microscopy (Leica, Wetzlar, Germany).

### Statistical analysis

Results from survival assays were analysed by two-way analysis of variance (ANOVA) with Tukey’s post hoc test for 95 % confidence intervals. For the qPCR results the relative fold induction (ΔΔCt) of the target gene compared to a normalised control sample was calculated and analysed by two-tailed Mann-Whitney U test. Statistical analyses were performed using GraphPad Prism version 9.1.0 for Windows (GraphPad Software, San Diego, California, USA). In figures, ns stands for p> 0.05, * p < 0.05, ** p < 0.01, *** p < 0.001, **** p < 0.0001. Error bars in figures specify SEM from the indicated number of independent experimental repeats. Experiments were performed indicated number of times (n ≥ 3) and statistics were calculated for each individual experiment.

## Acknowledgements

We thank Mads Gyrd Hansen for the M1-TUBE construct and Diana Toivola for Caco2 cells. We thank Pascal Meier, François Leulier, Bloomington *Drosophila* Stock Center (NIH P40OD018537), KYOTO Stock Center (DGRC) in Kyoto Institute of Technology and Vienna *Drosophila* Resource Center (VDRC) for fly stocks used in this study. Freddy Suarez Rodriguez and Cecilia Sahlgren are acknowledged for assistance with experiments involving mechanical stress. Emmy Himmelroos and Marc Sundén are acknowledged for technical assistance. The Academy of Finland (#321850), Sigrid Juselius Foundation, Åbo Akademi University Foundation, Victoriastiftelsen, Svenska Kulturfonden, Orion Research Foundation sr, and Turku Doctoral Network in Molecular Biosciences are acknowledged for financial support. This research was supported by the Center of Excellence in Cellular Mechanostasis at Åbo Akademi University and the InFLAMES Flagship Programme of the Academy of Finland (# 337530).

## Author contributions

ALA designed and executed most of the experiments, data analysis and writing of the manuscript. NT planned and performed the mechanical stress experiments in fly and cell samples. GMC planned and performed the experiments in cell samples. JK and PK planned and assembled the MiniHypoxy platform. MB contributed to the design of the experiments. AM contributed to the design of the experiments, writing and data analysis of this manuscript.

## Conflict of interest

The authors declare no conflict of interest.

## Notes

### Competing Interest Statement

The authors have declared no competing interest.

## References

1 Hershko A & Ciechanover A (1998) The ubiquitin system. Annu Rev Biochem 67, 425–479.

2 Dikic I, Wakatsuki S & Walters KJ (2009) Ubiquitin-binding domains from structures to functions. Nat Rev Mol Cell Biol 10, 659–671.

3 Swatek KN & Komander D (2016) Ubiquitin modifications. Cell Res 26, 399–422.

4 Komander D & Rape M (2012) The Ubiquitin Code. Annu Rev Biochem 81, 203–229.

5 Kirisako T, Kamei K, Murata S, Kato M, Fukumoto H, Kanie M, Sano S, Tokunaga F, Tanaka K & Iwai K (2006) A ubiquitin ligase complex assembles linear polyubiquitin chains. EMBO J 25, 4877–4887.

6 Gerlach B, Cordier SM, Schmukle AC, Emmerich CH, Rieser E, Haas TL, Webb AI, Rickard JA, Anderton H, Wong WW, Nachbur U, Gangoda L, Warnken U, Purcell AW, Silke J & Walczak H (2011) Linear ubiquitination prevents inflammation and regulates immune signalling. Nature 471, 591–596.

7 Ikeda F, Deribe YL, Skånland SS, Stieglitz B, Grabbe C, Franz-Wachtel M, Van Wijk SJL, Goswami P, Nagy V, Terzic J, Tokunaga F, Androulidaki A, Nakagawa T, Pasparakis M, Iwai K, Sundberg JP, Schaefer L, Rittinger K, MacEk B & Dikic I (2011) SHARPIN forms a linear ubiquitin ligase complex regulating NF-κB activity and apoptosis. Nature 471, 637–641.

8 Tokunaga F, Nakagawa T, Nakahara M, Saeki Y, Taniguchi M, Sakata SI, Tanaka K, Nakano H & Iwai K (2011) SHARPIN is a component of the NF-κB-activating linear ubiquitin chain assembly complex. Nature 471, 633–636.

9 Shimizu Y, Taraborrelli L & Walczak H (2015) Linear ubiquitination in immunity. Immunol Rev 266, 190–207.

10 Jahan AS, Elbæk CR & Damgaard RB (2021) Met1-linked ubiquitin signalling in health and disease: inflammation, immunity, cancer, and beyond. Cell Death Differ 28, 473–492.

11 Fiil BK & Gyrd-Hansen M (2014) Met1-linked ubiquitination in immune signalling. FEBS J 281, 4337–4350.

12 Aalto AL, Mohan AK, Schwintzer L, Kupka S, Kietz C, Walczak H, Broemer M & Meinander A (2019) M1-linked ubiquitination by LUBEL is required for inflammatory responses to oral infection in Drosophila. Cell Death Differ 26, 860–876.

13 Ferrandon D, Imler JL, Hetru C & Hoffmann JA (2007) The Drosophila systemic immune response: Sensing and signalling during bacterial and fungal infections. Nat Rev Immunol 7, 862–874.

14 Charroux B & Royet J (2010) Drosophila immune response: From systemic antimicrobial peptide production in fat body cells to local defense in the intestinal tract. Fly (Austin) 4, 40–47.

15 Lemaitre B & Hoffmann J (2007) The Host Defense of Drosophila melanogaster. Annu Rev Immunol 25, 697–743.

16 Gottar M, Gobert V, Michel T, Belvin M, Duyk G, Hoffmann JA, Ferrandon D & Royet J (2002) The Drosophila immune response against Gram-negative bacteria is mediated by a peptidoglycan recognition protein. Nature 416, 640–644.

17 Paquette N, Broemer M, Aggarwal K, Chen L, Husson M, Ertürk-Hasdemir D, Reichhart J-MM, Meier P, Silverman N, Erturk-Hasdemir D, Reichhart J-MM, Meier P & Silverman N (2010) Caspase-mediated cleavage, IAP binding, and ubiquitination: linking three mechanisms crucial for Drosophila NF-kappaB signaling. Mol Cell 37, 172–182.

18 Choe KM, Lee H & Anderson K V. (2005) Drosophila peptidoglycan recognition protein LC (PGRP-LC) acts as a signal-transducing innate immune receptor. Proc Natl Acad Sci U S A 102, 1122–1126.

19 Rutschmann S, Jung AC, Zhou R, Silverman N, Hoffmann JA & Ferrandon D (2000) Role of Drosophila IKKγ in a Toll-independent antibacterial immune response. Nat Immunol 1, 342–347.

20 Lu Y, Wu LP & Anderson K V. (2001) The antibacterial arm of the Drosophila innate immune response requires an IκB kinase. Genes Dev 15, 104–110.

21 Silverman N, Zhou R, Stöven S, Pandey N, Hultmark D & Maniatis T (2000) A Drosophila IκB kinase complex required for relish cleavage and antibacterial immunity. Genes Dev 14, 2461–2471.

22 Ertürk-Hasdemir D, Broemer M, Leulier F, Lane WS, Paquette N, Hwang D, Kim C-H, Stöven S, Meier P & Silverman N (2009) Two roles for the Drosophila IKK complex in the activation of Relish and the induction of antimicrobial peptide genes. Proc Natl Acad Sci U S A 106, 9779–84.

23 Stöven S, Silverman N, Junell A, Hedengren-Olcott M, Erturk D, Engström Y, Maniatis T & Hultmark D (2003) Caspase-mediated processing of the drosophila NF-κB factor relish. Proc Natl Acad Sci U S A 100, 5991–5996.

24 Valanne S, Wang J-H & Rämet M (2011) The Drosophila Toll Signaling Pathway. J Immunol 186, 649–656.

25 Cohen Peleg Rider I, Voronov E & Dinarello CA (2017) Alarmins: Feel the Stress. J Immunol Ref 198, 1395–1402.

26 Asri RM, Salim E, Nainu F, Hori A & Kuraishi T (2019) Sterile induction of innate immunity in Drosophila melanogaster. Front Biosci - Landmark 24, 1390–1400.

27 Rubartelli A & Sitia R (2009) Stress as an intercellular signal: the emergence of stress-associated molecular patterns (SAMP). Antioxid Redox Signal 11, 2621–2629.

28 Metsälä O, Kreutzer J, Högel H, Miikkulainen P, Kallio P & Jaakkola PM (2018) Transportable system enabling multiple irradiation studies under simultaneous hypoxia in vitro. Radiat Oncol 13, 220.

29 Nambu JR, Chen W, Hu S & Crews ST (1996) The Drosophila melanogaster similar bHLH-PAS gene encodes a protein related to human hypoxia-inducible factor 1α and Drosophila single-minded. Gene 172, 249–254.

30 Lavista-Llanos S, Centanin L, Irisarri M, Russo DM, Gleadle JM, Bocca SN, Muzzopappa M, Ratcliffe PJ & Wappner P (2002) Control of the hypoxic response in Drosophila melanogaster by the basic helix-loop-helix PAS protein similar. Mol Cell Biol 22, 6842–53.

31 Dekanty A, Romero NM, Bertolin AP, Thomas MG, Leishman CC, Perez-Perri JI, Boccaccio GL & Wappner P (2010) Drosophila genome-wide RNAi screen identifies multiple regulators of HIF-dependent transcription in hypoxia. PLoS Genet 6, e1000994.

32 Liu G, Roy J & Johnson EA (2006) Identification and function of hypoxia-response genes in Drosophila melanogaster. Physiol Genomics 25.

33 Arquier N, Vigne P, Duplan E, Hsu T, Therond PP, Frelin C & D’Angelo G (2006) Analysis of the hypoxia-sensing pathway in *Drosophila melanogaster*. Biochem J 393, 471–480.

34 Bruick RK & McKnight SL (2001) A conserved family of prolyl-4-hydroxylases that modify HIF. Science (80-) 294, 1337–1340.

35 Bandarra D, Biddlestone J, Mudie S, Muller HA & Rocha S (2014) Hypoxia activates IKK-NF-κB and the immune response in Drosophila melanogaster. Biosci Rep 34.

36 Boutros M, Agaisse H & Perrimon N (2002) Sequential activation of signaling pathways during innate immune responses in Drosophila. Dev Cell 3, 711–722.

37 Buchon N, Broderick NA, Poidevin M, Pradervand S & Lemaitre B (2009) Drosophila Intestinal Response to Bacterial Infection: Activation of Host Defense and Stem Cell Proliferation. Cell Host Microbe 5, 200–211.

38 Tzou P, Ohresser S, Ferrandon D, Capovilla M, Reichhart J-M, Lemaitre B, Hoffmann JA & Imler J-L (2000) Tissue-Specific Inducible Expression of Antimicrobial Peptide Genes in Drosophila Surface Epithelia. Immunity 13, 737–748.

39 Davis MM & Engström Y (2012) Immune response in the barrier epithelia: Lessons from the fruit fly drosophila melanogaster. J Innate Immun 4, 273–283.

40 Ferrandon D, Jung AC, Criqui MC, Lemaitre B, Uttenweiler-Joseph S, Michaut L, Reichhart JM & Hoffmann JA (1998) A drosomycin-GFP reporter transgene reveals a local immune response in Drosophila that is not dependent on the Toll pathway. EMBO J 17, 1217–1227.

41 Akhouayri I, Turc C, Royet J & Charroux B (2011) Toll-8/Tollo negatively regulates antimicrobial response in the Drosophila respiratory epithelium. PLoS Pathog 7, e1002319.

42 Hedengren M, Åsling B, Dushay MS, Ando I, Ekengren S, Wihlborg M & Hultmark D (1999) Relish, a central factor in the control of humoral but not cellular immunity in Drosophila. Mol Cell 4, 827–837.

43 Hosamani R & Muralidhara (2013) Acute exposure of drosophila melanogaster to paraquat causes oxidative stress and mitochondrial dysfunction. Arch Insect Biochem Physiol 83, 25–40.

44 Bus JS & Gibson JE (1984) Paraquat: model for oxidant-initiated toxicity. Environ Health Perspect 55, 37–46.

45 Kenmoku H, Hori A, Kuraishi T & Kurata S (2017) A novel mode of induction of the humoral innate immune response in Drosophila larvae. DMM Dis Model Mech 10, 271–281.

46 Noad J, Von Der Malsburg A, Pathe C, Michel MA, Komander D & Randow F (2017) LUBAC-synthesized linear ubiquitin chains restrict cytosol-invading bacteria by activating autophagy and NF-κB. Nat Microbiol 2, 1–10.

47 van Well EM, Bader V, Patra M, Sánchez-Vicente A, Meschede J, Furthmann N, Schnack C, Blusch A, Longworth J, Petrasch-Parwez E, Mori K, Arzberger T, Trümbach D, Angersbach L, Showkat C, Sehr DA, Berlemann LA, Goldmann P, Clement AM, Behl C, Woerner AC, Saft C, Wurst W, Haass C, Ellrichmann G, Gold R, Dittmar G, Hipp MS, Hartl FU, Tatzelt J & Winklhofer KF (2019) A protein quality control pathway regulated by linear ubiquitination. EMBO J 38.

48 Schaeffer V, Akutsu M, Olma MH, Gomes LC, Kawasaki M & Dikic I (2014) Binding of OTULIN to the PUB Domain of HOIP Controls NF-κB Signaling. Mol Cell 54, 349–361.

49 Takiuchi T, Nakagawa T, Tamiya H, Fujita H, Sasaki Y, Saeki Y, Takeda H, Sawasaki T, Buchberger A, Kimura T & Iwai K (2014) Suppression of LUBAC-mediated linear ubiquitination by a specific interaction between LUBAC and the deubiquitinases CYLD and OTULIN. Genes to Cells 19, 254–272.

50 Asaoka T, Almagro J, Ehrhardt C, Tsai I, Schleiffer A, Deszcz L, Junttila S, Ringrose L, Mechtler K, Kavirayani A, Gyenesei A, Hofmann K, Duchek P, Rittinger K & Ikeda F (2016) Linear ubiquitination by LUBEL has a role in Drosophila heat stress response. EMBO Rep 17, 1624–1640.

51 Leulier F, Rodriguez A, Khush RS, Abrams JM & Lemaitre B (2000) The Drosophila caspase Dredd is required to resist Gram-negative bacterial infection. EMBO Rep 1, 353–358.

52 Välimäki H, Verho J, Kreutzer J, Kattipparambil Rajan D, Ryynänen T, Pekkanen-Mattila M, Ahola A, Tappura K, Kallio P & Lekkala J (2017) Fluorimetric oxygen sensor with an efficient optical read-out for in vitro cell models. Sensors Actuators, B Chem 249, 738–746.

53 Driessen R, Zhao F, Hofmann S, Bouten C, Sahlgren C & Stassen O (2020) Computational characterization of the dish-in-a-dish, a high yield culture platform for endothelial shear stress studies on the orbital shaker. Micromachines 11, 552.

54 Fiil BK, Damgaard RB, Wagner SA, Keusekotten K, Fritsch M, Bekker-Jensen S, Mailand N, Choudhary C, Komander D & Gyrd-Hansen M (2013) OTULIN Restricts Met1-Linked Ubiquitination to Control Innate Immune Signaling. Mol Cell 50, 818–830.

55 Keusekotten K, Elliott PR, Glockner L, Fiil BK, Damgaard RB, Kulathu Y, Wauer T, Hospenthal MK, Gyrd-Hansen M, Krappmann D, Hofmann K & Komander D (2013) OTULIN antagonizes LUBAC signaling by specifically hydrolyzing met1-linked polyubiquitin. Cell 153, 1312.

